# Exploring novel inducers of phage tail-like particle expression using a fluorescent reporter bacterium

**DOI:** 10.1101/2025.10.13.681979

**Authors:** Jordan Vacheron, Clara M. Heiman, Martina Crea, Alexandre Erbetta, Christoph Keel

**Author notes:** **Correspondence:** Jordan Vacheron, Christoph Keel. Nestlé, Route du Jorat 57, CH-1000 Lausanne, Switzerland. Nestlé, Chemin de Rive 5, CH-1350 Orbe, Switzerland.

## Abstract

R-tailocins are phage tail-like particles produced by diverse bacterial species, playing crucial roles in interbacterial competition and microbial community dynamics. While their production is typically triggered by DNA damage through the bacterial SOS response, the full range of potential inducers remains largely unexplored. This study aimed at identifying novel chemical compounds capable of triggering R-tailocin gene expression. A fluorescent reporter bacterial strain was employed to monitor R-tailocin gene cluster activity, and over 1,500 compounds were screened using Biolog Phenotype MicroArray plates. This screen identified several previously unreported inducers of R-tailocin expression. Among the strongest inducers identified were pipemidic acid and hydroxyurea, although their induction levels did not exceed those observed with the commonly used inducer mitomycin C. The identified compounds exhibited distinct temporal induction patterns, suggesting variation in their underlying regulatory mechanisms. Several also displayed clear dose-response patterns, providing further insights into their potency and potential modes of action. Notably, R-tailocin induction did not necessarily lead to complete population lysis, as a subset of cells consistently survived the induction process. Collectively, these findings expand the current understanding of R-tailocin induction and production in bacterial populations, uncovering novel chemical triggers and revealing diverse expression dynamics. These insights offer potential implications for microbial community ecology and the development of targeted antimicrobial strategies.

## Introduction

R-tailocins are phage tail-like particles produced by many bacterial species, including members of the genus *Pseudomonas* (Dorosky *et al*. 2017; Vacheron, Heiman and Keel 2021; Heiman, Vacheron and Keel 2023). These high-molecular-weight protein complexes resemble the tails of bacteriophages and are encoded in gene clusters within bacterial genomes (Heiman, Vacheron and Keel 2023). R-tailocins are highly specialized structures that play crucial roles in interbacterial competition and ecology, influencing microbial community dynamics by specifically killing closely related strains (Vacheron, Heiman and Keel 2021; Heiman, Vacheron and Keel 2023). Recent advances in genomics and molecular biology have enabled the identification and characterization of R-tailocin and more broadly extracellular contractile injections systems (eCIS) gene clusters in various bacterial species (Geller *et al*. 2021; Heiman, Vacheron and Keel 2023). Although, the genetic clusters encoding these particles often carry the same basic gene set (structural components, lysis cassettes, and regulatory elements), the organization and regulation of these clusters can vary between species and even between strains of the same species, reflecting the diversity and specificity of R-tailocin production (Ghequire and De Mot 2015; Scholl 2017; Patz *et al*. 2019; Heiman, Vacheron and Keel 2023; Heiman *et al*. 2025).

The production of R-tailocins is induced under stresses that cause DNA damage, which activates the SOS response (Matsui *et al*. 1993; Heiman, Vacheron and Keel 2023; Heiman *et al*. 2025). Mitomycin C, a potent DNA cross-linking agent, is commonly used in studies to induce these phage tail-like particles and phages more broadly (Otsuji *et al*. 1959; Penterman, Singh and Walker 2014; Heiman, Vacheron and Keel 2023). Although other DNA damaging factors such as ultraviolet rays or ciprofloxacin have been used to elicit R-tailocin expression, mitomycin C has been shown to be the most effective in triggering the release of R-tailocins and prophages in various bacterial species, making it a fundamental tool for investigating induction mechanisms. However, the full range of environmental and chemical signals that can potentially trigger R-tailocin production remains largely unexplored. Identifying these inducers and understanding their induction mechanisms is essential for further elucidating the ecological role of R-tailocins and their potential applications in microbiome engineering and antimicrobial therapy.

*Pseudomonas protegens* CHA0 is a root-colonizing bacterium that is widely used as an environmental model strain for the *Pseudomonas* genus. The genome of this strain harbors three different phage-related clusters including one that encodes two R-tailocins (Vacheron, Heiman and Keel 2021). We used a fluorescent reporter strain of *P. protegens* CHA0 that allows for real-time monitoring of R-tailocin gene expression (Vacheron, Heiman and Keel 2021; Heiman *et al*. 2025) to systematically explore the induction of R-tailocin gene clusters. By using the Biolog™ Phenotype MicroArray plates (PM1 to PM20), which contain a wide range of chemical compounds and growth conditions, we aimed at identifying novel inducers of R-tailocin expression beyond the commonly used mitomycin C. This approach provided a powerful tool for screening potential inducers and studying the regulation of R-tailocin production. Our study reveals new insights into the environmental and chemical regulation of R-tailocin production, emphasizing the influence of common microbiological growth conditions.

## Results and discussion

### Identification of novel inducers of R-tailocin expression

To identify novel chemical triggers of R-tailocin expression, we screened over 1,500 compounds by following the growth and fluorescence of a fluorescent R-tailocin reporter strain, CHA0 pOT1e-P_*hol*_-*egfp*, in Biolog Phenotype MicroArray plates (PM1 to PM20) (**Figure 1, supplementary Figures S1-S7**). Several previously unrecognized inducers were detected, with notable candidates including nitrogen-containing heterocycles, such as pipemidic acid derivatives and urea-based compounds (**Figure 1**). These findings align with previous studies suggesting that nitrogen metabolism and stress responses may play a role in phage-related gene expression (Nilsson *et al*. 2022). Pipemidic acid is a quinolone that inhibits DNA synthesis by targeting class II topoisomerases such as the DNA gyrase and the topoisomerase I, resulting in DNA damage and the activation of the bacterial SOS. This stress response, in turn, induces the expression of prophages and tailocins (Shimizu *et al*. 1975; Mitscher 2005). Similarly, urea-based compounds interfere with DNA integrity and gene expression at multiple levels, including RNA and protein unfolding (Raghunathan, Jaganade and Priyakumar 2020). At the DNA level, urea disrupts intramolecular interactions, leading to strand denaturation and inhibition of DNA replication (Raghunathan, Jaganade and Priyakumar 2020). Thus, both pipemidic acid and urea-based compounds compromise DNA stability, replication or repair in some way, ultimately triggering R-tailocin via activation of the bacterial SOS system.

**Figure 1:**
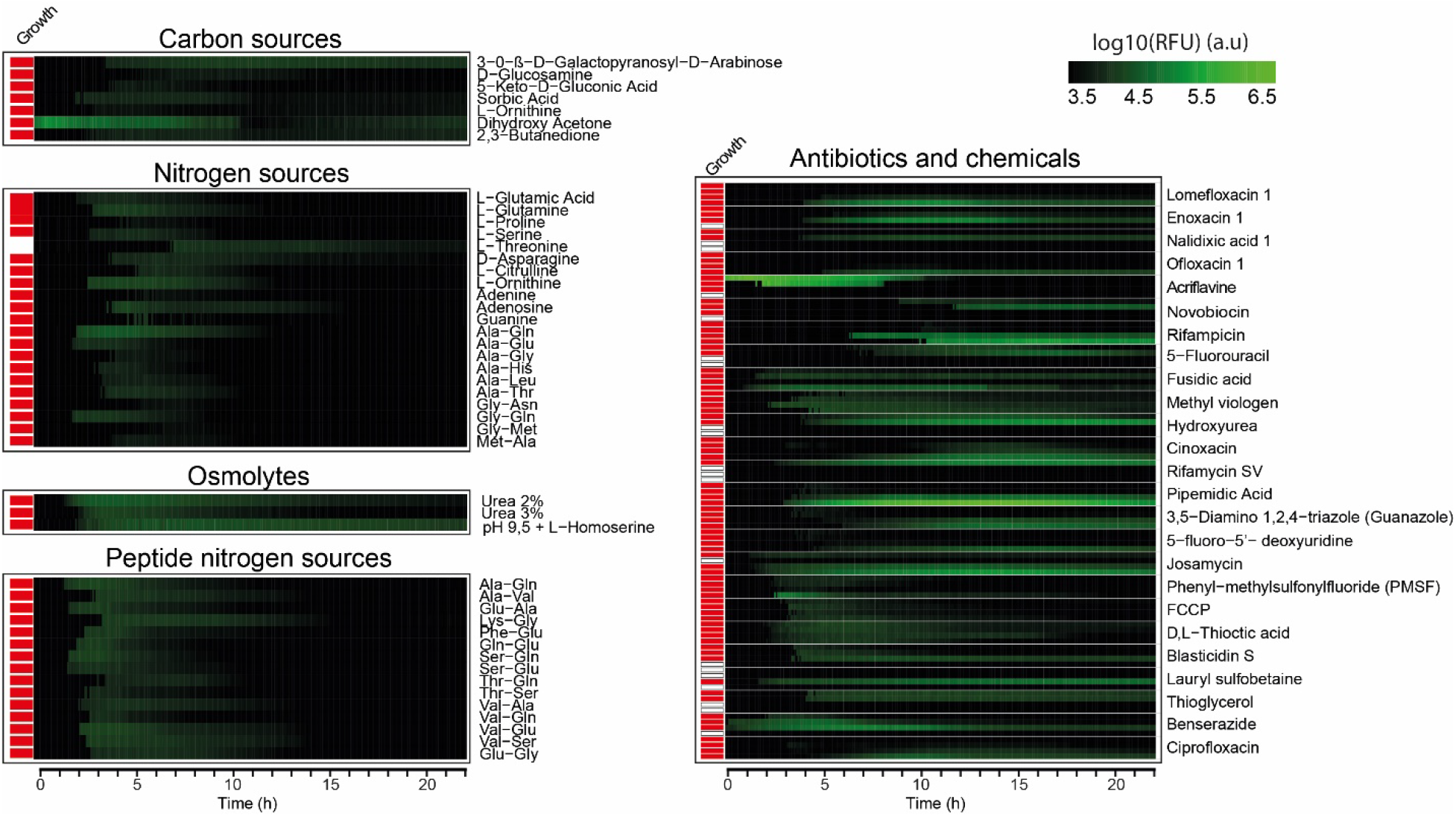
Induction patterns of R-tailocin expression by different compounds. Heatmaps showing fluorescence intensity over time for identified R-tailocin inducers. Red squares indicate bacterial growth; white squares indicate no growth. Complete datasets are provided in Supplementary Figures S1–S7 and in the analyzed dataset deposited on Zenodo.

Interestingly, several carbon and nitrogen sources commonly used in microbiology exhibited mild R-tailocin induction (**Figure 1**). Indeed, sugars such as 3-0-β-D-galactopyranosyl-D-arabinose (a rare sugar found in plants), D-glucosamine (an amino-sugar widely found in nature), dihydroxy acetone or 2,3-butanedione (volatile organic compound and byproduct of fermentation) appear to induce R-tailocin expression. Additionally, nitrogen sources such as Alanine-Glutamine, L-ornithine, adenosine (that are commonly used in the laboratory) seem to induce the expression of the R-tailocin gene cluster. Interestingly, L-glutamine induces tailocin expression and has been shown to influence the spatial colonization of root-associated microbes via root leakage (Tsai *et al*. 2025). These observations suggest that standard culture conditions may inadvertently influence R-tailocin-related dynamics and should be considered when interpreting microbial competition assays.

### Induction of the R-tailocin gene cluster expression is independent of the mode of action of a compound

Among the chemical inducers identified, the most potent belonged to antibiotic and antimicrobial compound classes (**Figure 1**). Besides mitomycin C, pipemidic acid and urea, several other chemical and antibiotic compounds were found to induce R-tailocin expression. Most of these candidate inducers affect DNA transcription, replication and/or repair. Acriflavine, originally used as a surface antiseptic during the World War I against sleeping sickness, is also an effective antibacterial chemotherapy agent that intercalates DNA (Browning 1943; Nakamura and Suganuma 1972; Piorecka, Kurjata and Stanczyk 2022). It strongly induced R-tailocin expression (**Figure 1**). Similarly, hydroxyurea also led to R-tailocin expression. The compound, which is known to activate the SOS response in *Bacillus subtilis* and *Escherichia coli*, reduces the pool of deoxyribonucleoside triphosphate (dNTP) by inhibiting ribonucleotide reductase (Rosenkranz *et al*. 1967; Sinha and Snustad 1972; Wozniak and Simmons). Similarly to pipemidic acid, lomefloxacin, an antibiotic that interferes with DNA gyrase and topoisomerase IV (Piddock, Hall and Wise 1990), also induced R-tailocin expression (**Figure 1**). This fluoroquinolone has previously been shown to increase the expression of tailocins (R/F-pyocins) in *Pseudomonas aeruginosa* PA14 (Franke *et al*. 2021). At the level of DNA transcription, rifampicin, an antibiotic targeting DNA transcription by binding to the RNA polymerase (Wehrli 1983), induced R-tailocin expression as well (**Figure 1**). Finally, at the level of translation, josamycin, which inhibits protein synthesis by binding to the 50S subunit of the ribosome (Tenson, Lovmar and Ehrenberg 2003), also acted as R-tailocin inducer (**Figure 1**).

In summary, amongst all the compounds tested, all those known to damage DNA or interfere with DNA-related processes, except for proflavine, were found to induce R-tailocin expression. These findings suggest that activation of the R-tailocin gene cluster is broadly associated with cellular stress, particularly that which affects DNA integrity, but is not strictly dependent on a specific mode of action.

A principal component analysis (PCA) was performed on the chemical and antibiotic compounds to assess whether distinct groups can be discerned (**Figure 2**). The analysis revealed that all tested compounds triggered R-tailocin expression with comparable intensities (**Figure 2**). This observation is in line with previous findings, as R-tailocins, similar to their prophage ancestors, are expressed in response to DNA damage, which activates RecA and the bacterial SOS system (Matsui *et al*. 1993; Penterman, Singh and Walker 2014; Heiman, Vacheron and Keel 2023; Heiman *et al*. 2025).

**Figure 2:**
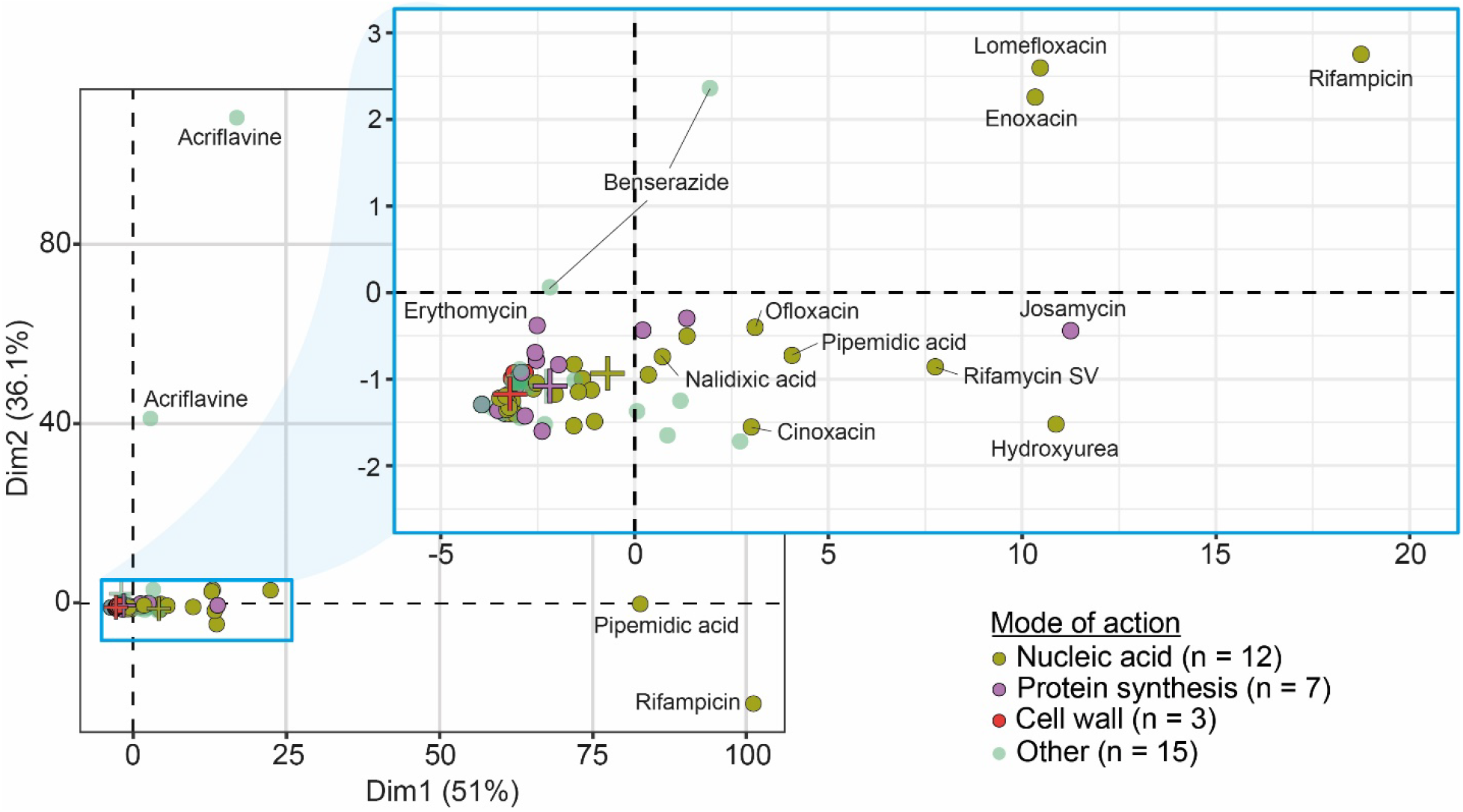
Principal component analysis (PCA) of R-tailocin-inducing compounds. PCA visualization of compounds inducing R-tailocin expression, grouped by their mode of action. Clustering indicates distinct chemical classes of inducers. Colored crosses represent the barycenters of each group’s distribution.

### Selection of the best inducers and their dose-responses effects

From the full screening dataset, we selected a subset of compounds showing the strongest and most consistent induction of R-tailocin expression. These included hydroxyurea, pipemidic acid, acriflavine, rifampicin, lomefloxacin, enoxacin, ofloxacin, nalidixic acid, erythromycin, novobiocin, chloramphenicol, and colistin (**Figure 3**). These molecules were selected because they not only belong to pharmacologically diverse classes, such as DNA-interacting agents, transcription inhibitors, and protein synthesis inhibitors, but also exhibited clear and strong dose-dependent increases in fluorescence, indicating a robust activation of the R-tailocin gene cluster. Such patterns suggest that R-tailocin induction may be linked to general cellular stress responses rather than a specific molecular target. However, due to the limited and undisclosed concentration range provided by the Biolog system, precise activation thresholds could not be determined. While mitomycin C remained the most effective inducer, our data highlight novel compounds with potential applications in microbial ecology and antimicrobial strategy development.

**Figure 3.**
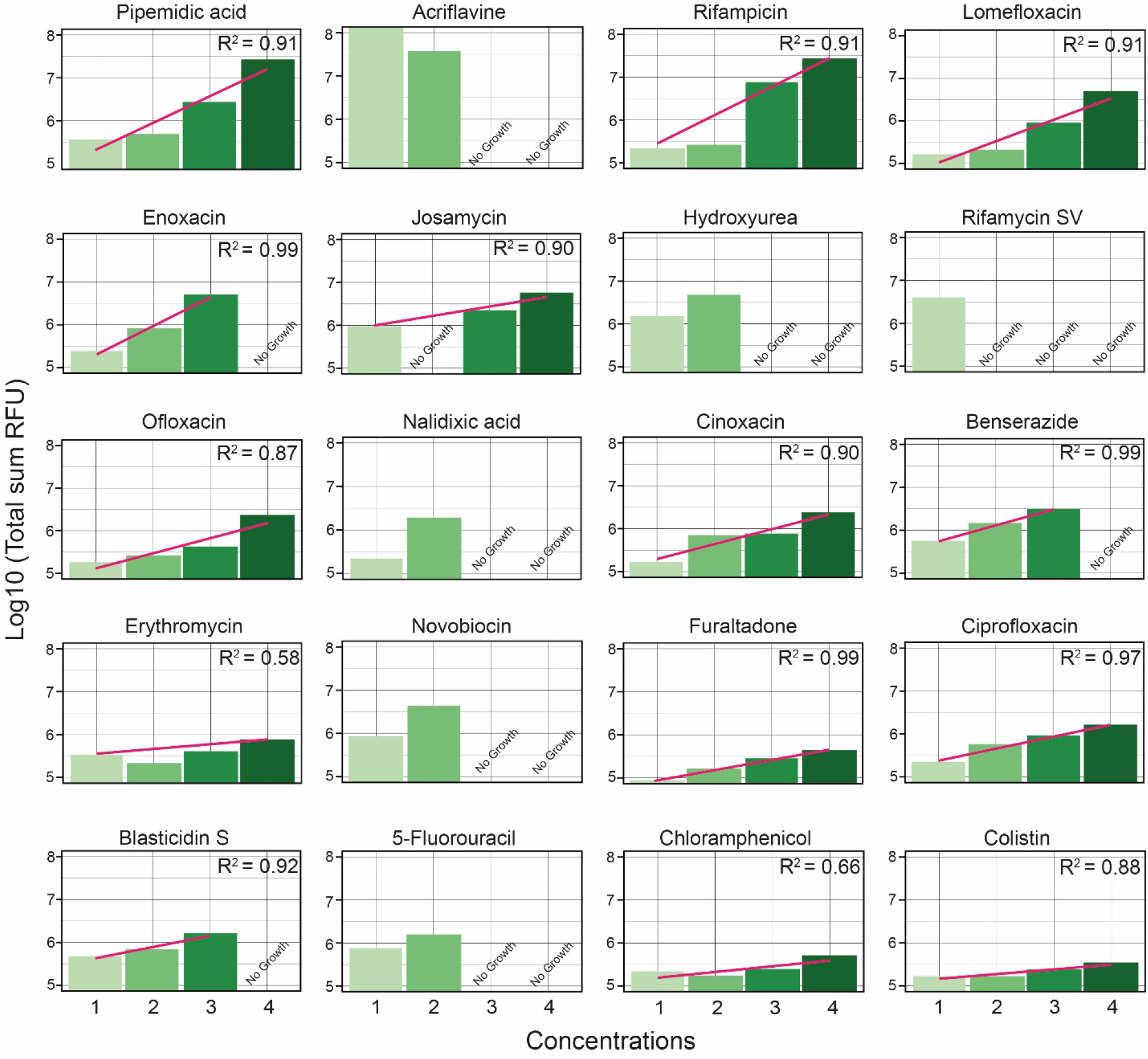
Dose-response relationships of selected inducers. The sum of the Log10-transformed fluorescence (RFU) plotted against increasing compound concentrations. Red lines represent linear regression fits; R^2^ values indicate correlation strength. Absence of bacterial growth is indicated with the label “No Growth”.

### Validation with R-tailocin extraction and semi quantification

To confirm that the transcriptional induction observed in the reporter assay translates into actual R-tailocin production, we selected a subset of compounds representing distinct chemical classes and strong dose-dependent responses in fluorescence assays, **acriflavine, rifampicin, pipemidic acid, and hydroxyurea**, for experimental validation (Figure 4). R-tailocin preparations obtained from *P. protegens* CHA0 cultures treated with acriflavine, rifampicin, pipemidic acid, or hydroxyurea were compared to those obtained after mitomycin C treatment, which served as a positive control. Serial dilutions of the extracts were spotted onto soft agar overlays containing *P. protegens* Pf-5, a strain sensitive to the CHA0-derived R-tailocins. As expected, MMC treatment resulted in strong and clear lytic zones, confirming efficient induction of R-tailocin production. Pipemidic acid treatment yielded detectable lytic activity, albeit weaker than that induced by MMC. Hydroxyurea induced detectable lysis as well, but the activity was weaker than that observed with MMC and pipemidic acid. Acriflavine produced weak but visible activity, limited to the first dilution spot. In contrast, rifampicin appeared to produce no detectable activity in these assays; however, this apparent absence of activity may be attributed to technical limitations in interpreting the transcriptional reporter data. As the reporter is based on GFP, and rifampicin solutions display a yellow– orange coloration depending on concentration, interference with fluorescence detection remains possible despite the use of an appropriate control. No activity was observed in extracts from the *P. protegens* CHA0 Δ4 strain (deleted for all prophages and R-tailocin loci) in any condition, confirming that the observed killing was attributable to R-tailocin particles.

**Figure 4.**
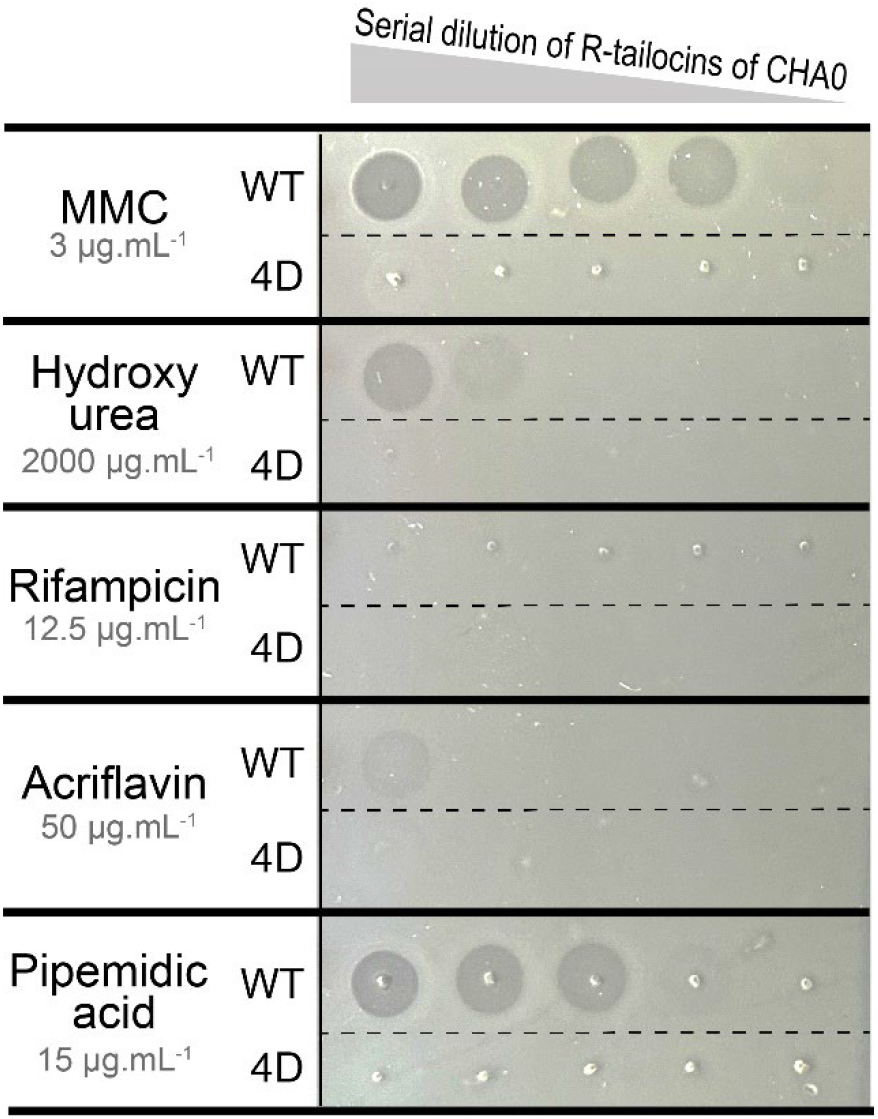
Validation of R-tailocin activity after induction with selected compounds. R-tailocins were induced in *Pseudomonas protegens* CHA0 using different chemical inducers at the concentrations indicated below each compound name. Serial dilutions of purified R-tailocins were spotted on soft agar overlays seeded with the sensitive strain *P. protegens* Pf-5. “WT” corresponds to R-tailocins extracted from the CHA0 wild type, and “4D” to extracts from a CHA0 derivative deleted for all prophages and R-tailocin loci. MMC, mitomycin C.

## Conclusion

In this study, we systematically screened a diverse set of chemical compounds to identify novel inducers of R-tailocin production. While several compounds had previously been associated with the triggering of the release of phages and phage tail-like particles, others, such as pipemidic acids and urea derivatives, emerged as newly identified activators. Notably, our results suggest that commonly used microbiological culture conditions may unintentionally trigger R-tailocin expression, underscoring the importance of careful experimental design in microbial interaction studies. Together, these findings enhance our understanding of R-tailocin regulation and open new avenues for the targeted manipulation of bacterial populations.

## Material and methods

### Bacterial strains and culture conditions

*Pseudomonas protegens* CHA0 wild type (WT), the Δ4 derivative (deleted for all prophages and R-tailocin loci), the transcriptional reporter strain for R-tailocin gene cluster expression, *Pseudomonas protegens* CHA0 pOT1e-P_*hol*_-*egfp* (Heiman *et al*. 2022, 2025), and the CHA0 R-tailocin sensitive strain *P. protegens* Pf-5 (Vacheron, Heiman and Keel 2021) were used in the assays (**Suppl. Table S1)**. Strains were routinely cultured at 25 °C on nutrient agar (NA) and in nutrient yeast broth (NYB), supplemented with gentamicin (10 µg mL^-1^) where required.

### Screening of potential R-tailocin expression inducers

Biolog™ Phenotype Microarray (PM) plates 1 to 20 (Biolog Inc. USA) were used to screen for inducers of the R-tailocin gene cluster of CHA0. Overnight cultures of the R-tailocin gene expression reporter CHA0 pOT1e-P_*hol*_-*egfp* were restarted into fresh NYB (1:100) supplemented with gentamycin (10 µg mL^-1^). When cells reached exponential growth phase i.e., at an optical density (OD_600nm_) of 0.4-0.6, the bacterial cells were washed with Minimal Medium (MM – 0,1% [w/v] NH_4_Cl; 3,49% [w/v] Na_2_HPO_4_.2H_2_O; 2,77% [w/v] KH_2_PO_4_; supplemented when needed with 80 mM of glucose) and concentrated to obtain a bacterial cell suspension adjusted to OD_600nm_ = 2 in MM. For the PM plates related to carbon utilization (#1 and #2), the bacterial cells were resuspended in MM without carbon source. For the PM plates related to nitrogen (#3, #6, #7 and #8) or phosphorus and sulfur (#4) utilization, the bacterial cells were resuspended in MM without nitrogen source or phosphorus source respectively. Aliquots of 95 μL of the respective MM preparation were added in each well of the 96-wells Biolog™ PM plates. The OD_600nm_ and the GFP fluorescence (excitation 479 nm, emission 520 nm) were monitored for 30 min every 5 min using a BioTek Synergy H1 plate reader (BioTek Instruments Inc., Winooski, VT, USA) to obtain blank values for each compound of the Biolog™ PM plates. Then, 5 μL of the bacterial suspension were added to the 95 µL medium already filled in the wells, leading to a final load of approximatively 10^6^ CFU mL^-1^ (OD_600nm_ = 0.1). The OD_600nm_ and the GFP fluorescence were monitored for 22 h every 5 min using the BioTek Synergy H1 plate reader.

### Data analysis of inducer screening

Data from the BioTek Synergy H1 plate reader were processed to calculate the reporter fluorescence response for each well. Raw OD_600_ and GFP fluorescence values were corrected by subtracting the corresponding blank values, which were obtained from wells containing medium and compounds without bacteria. To ensure robust data analysis, several filtering criteria were applied: First, bacterial growth in each well was assessed by comparing the OD_600_ at the final time point to the OD_600_ at the initial time point. Growth was considered to have occurred if the final OD_600_ value was at least twice the initial the initial one; wells that did not meet this criterion got their values set to 0. Next, fluorescence values were adjusted to account for background noise. Negative fluorescence values after blank correction were replaced with zero. Additionally, fluorescence values that were lower than the maximum fluorescence observed in the negative control wells (containing no inducer) were also set to zero. Then, the relative fluorescence unit (RFU) was calculated by dividing the corrected GFP fluorescence by the OD_600_ at each time point. If the OD_600_ was zero, RFU values were set to zero to avoid errors in the calculation. The RFU values were then log-transformed using log_10_ to normalize the data. In cases where RFU values were zero, their log-transformed value was also set to zero to avoid undefined results. Finally, the processed data were visualized using heatmaps to highlight the GFP induction pattern across compounds. To identify patterns in R-tailocin induction, Principal Component Analysis (PCA) was performed on fluorescence data from compounds exhibiting induction responses. Data were log-transformed and scaled before analysis. Data were analyzed using R studio version 4.3.1.

### R-tailocin extraction and sensitivity tests

*P. protegens* CHA0 WT and the Δ4 derivative (deleted for all prophages and R-tailocin loci) were used for R-tailocin induction assays. Overnight cultures grown in NYB were diluted 1:100 into fresh NYB and incubated for approximately 3 h until reaching the exponential growth stage. Cultures were then treated with either mitomycin C (3 µg·mL^−1^), hydroxyurea (2000 µg·mL^−1^), rifampicin (12.5 µg·mL^−1^), acriflavine (50 µg·mL^−1^), or pipemidic acid (15 µg·mL^−1^). Following overnight incubation, cells were pelleted by centrifugation, and the supernatant was filtered through a 0.2 µm pore-size membrane. R-tailocins were precipitated by adding polyethylene glycol (PEG 8000, 10% w/v) and incubating the mixture overnight at 4 °C. Precipitated particles were collected by centrifugation at maximum speed and resuspended in SM buffer (200 mM NaCl_2_; 10 mM MgSO_4_; 50 mM Tris-HCl; adjusted to pH 7.5), followed by gentle shaking at 4 °C overnight to ensure complete solubilization.

Semi-quantitative assessment of R-tailocin activity was performed by spotting serial dilutions of the purified preparations onto bacterial lawns. Briefly, 3 µL of each R-tailocin dilution were spotted onto NA plates overlaid with *P. protegens* Pf-5, a strain sensitive to the R-tailocin #1 of CHA0 (Vacheron, Heiman and Keel 2021). Plates were incubated overnight at 25 °C, after which clear halos of growth inhibition were observed and recorded.

## Acknowledgements

We thank Jeremi Cardon for technical assistance. We thank Nazife Beqa, Latifa Labidi and Nelson Galvao for their precious help preparing media and with material maintenance.

## Data availability

Dataset and raw data were deposited on Zenodo (10.5281/zenodo.17339175).

