## Supplementary material for "Exploring novel inducers of phage tail-like particle expression using a fluorescent reporter bacterium": Suppl. information

Jordan Vacheron[1#], Clara M. Heiman[1], Martina Crea[1*], Alexandre Erbetta[1$], and Christoph Keel[1#]
[1]Department of Fundamental Microbiology, University of Lausanne, Lausanne, Switzerland

[*]**Current address:** Nestlé, Route du Jorat 57, CH-1000 Lausanne, Switzerland.
[$]**Current address:** Nestlé, Chemin de Rive 5, CH-1350 Orbe, Switzerland

### Content:

**Supplementary Table 1**: Bacterial strains and plasmids used in this study.

**Figure S1:** Monitoring reporter fusion induction of R-tailocin expression across different carbon sources.
**Figure S2:** Monitoring reporter fusion induction of R-tailocin expression across different nitrogen sources (PM3) and sulfur- and phosphorus-based compounds (PM4).
**Figure S3:** Monitoring reporter fusion induction of R-tailocin expression in different biosynthetic pathways (PM5) and under different peptide compositions (PM6).
**Figure S4:** Monitoring reporter fusion induction of R-tailocin expression under different peptide compositions (PM7 and PM8).
**Figure S5:** Monitoring reporter fusion induction of R-tailocin expression under different osmotic stress (PM9) and pH (PM10) conditions.
**Figure S6:** Monitoring reporter fusion induction of R-tailocin expression under different chemical stress (1/2).
**Figure S7:** Monitoring reporter fusion induction of R-tailocin expression under different chemical stress (2/2).

**Supplementary Table 1**: Bacterial strains and plasmids used in this study.

| Strain / plasmid names | Genotype or relevant characteristics[1] | References |
|---|---|---|
| ***Pseudomonas protegens*** | | |
| CHA0 | *P. protegens* type strain; wild type; genome accession no. LS999205.1 | (Stutz 1986; Smits *et al.* 2019) |
| 4D (Δtail1Δtail2ΔmyoΔsiph) | CHA0 with the deletion of the sheath and tube genes of the tailocins #1 and #2 and of the Myoviridae and Siphoviridae prophages | (Vacheron, Heiman and Keel 2021) |
| Pf-5 | *P. protegens* sensitive to the R-tailocin #1 produced by CHA0 | (Vacheron, Heiman and Keel 2021) |
| **Plasmids** | | |
| pOT1e-$P_{hol}$-*egfp* | $P_{hol}$-*egfp* transcriptional fusion in pOT1e; used for monitoring R-tailocin gene cluster expression in CHA0; $Gm^R$ | (Heiman *et al.* 2025) |

[1] $Gm^R$, gentamycin resistance.

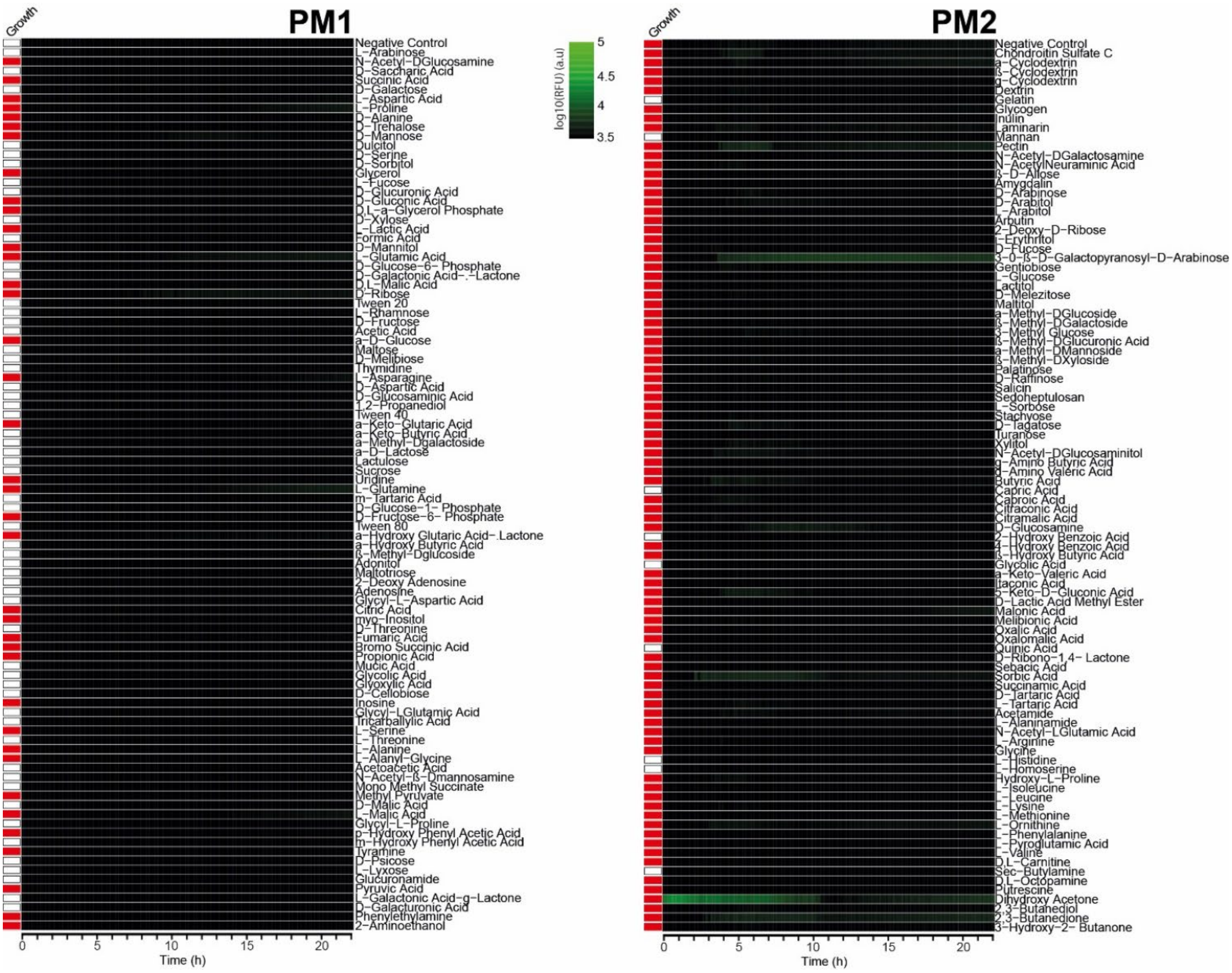


**Figure S1: Monitoring reporter fusion induction of R-tailocin expression across different carbon sources.**

Cells of *P. protegens* CHA0 carrying the R-tailocin reporter fusion were inoculated into Biolog PM1 and PM2 plates, each of which contain a distinct carbon source in every well. The inoculation medium consisted of Minimal Medium (MM) without any added carbon source (no glucose), so that the carbon compounds provided in the Biolog plates served as the sole available carbon sources for bacterial growth. RFU (Relative Fluorescence Unit) represents the fluorescence signal normalized by optical density. Red squares indicate bacterial growth, while white squares indicate the absence of bacterial growth.

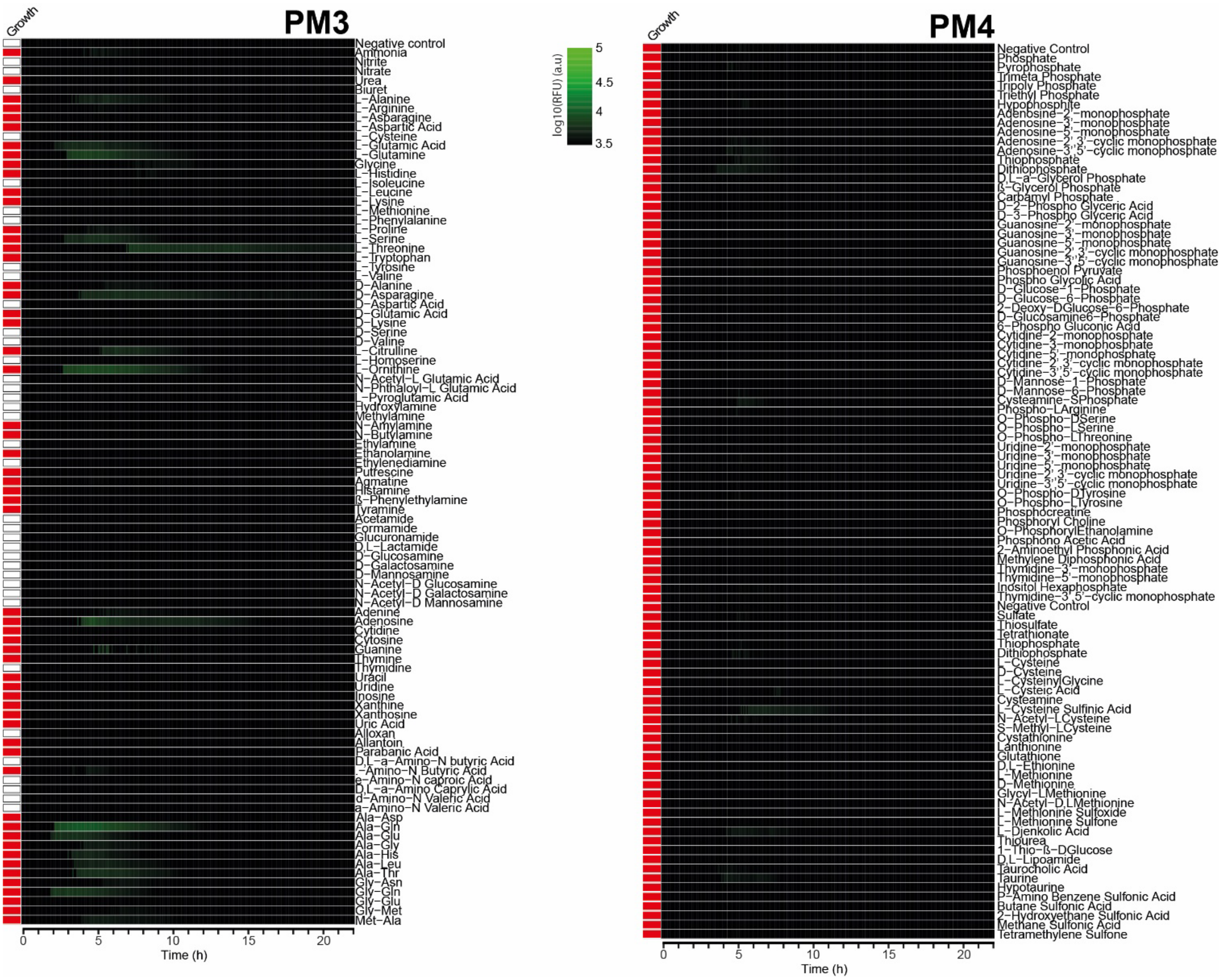


**Figure S2: Monitoring reporter fusion induction of R-tailocin expression across different nitrogen sources (PM3) and sulfur- and phosphorus-based compounds (PM4).**

Cells of *P. protegens* CHA0 carrying the R-tailocin reporter fusion were inoculated into Biolog PM3 and PM4 plates, each of which contains a distinct nitrogen source (PM3) and distinct phosphorus and sulfur sources (PM4) in every well. The inoculation medium consisted of Minimal Medium (MM) without any added nitrogen sources for PM3 and without phosphorus source for PM4. RFU (Relative Fluorescence Unit) represents the fluorescence signal normalized by optical density. Red squares indicate bacterial growth, while white squares indicate the absence of bacterial growth.

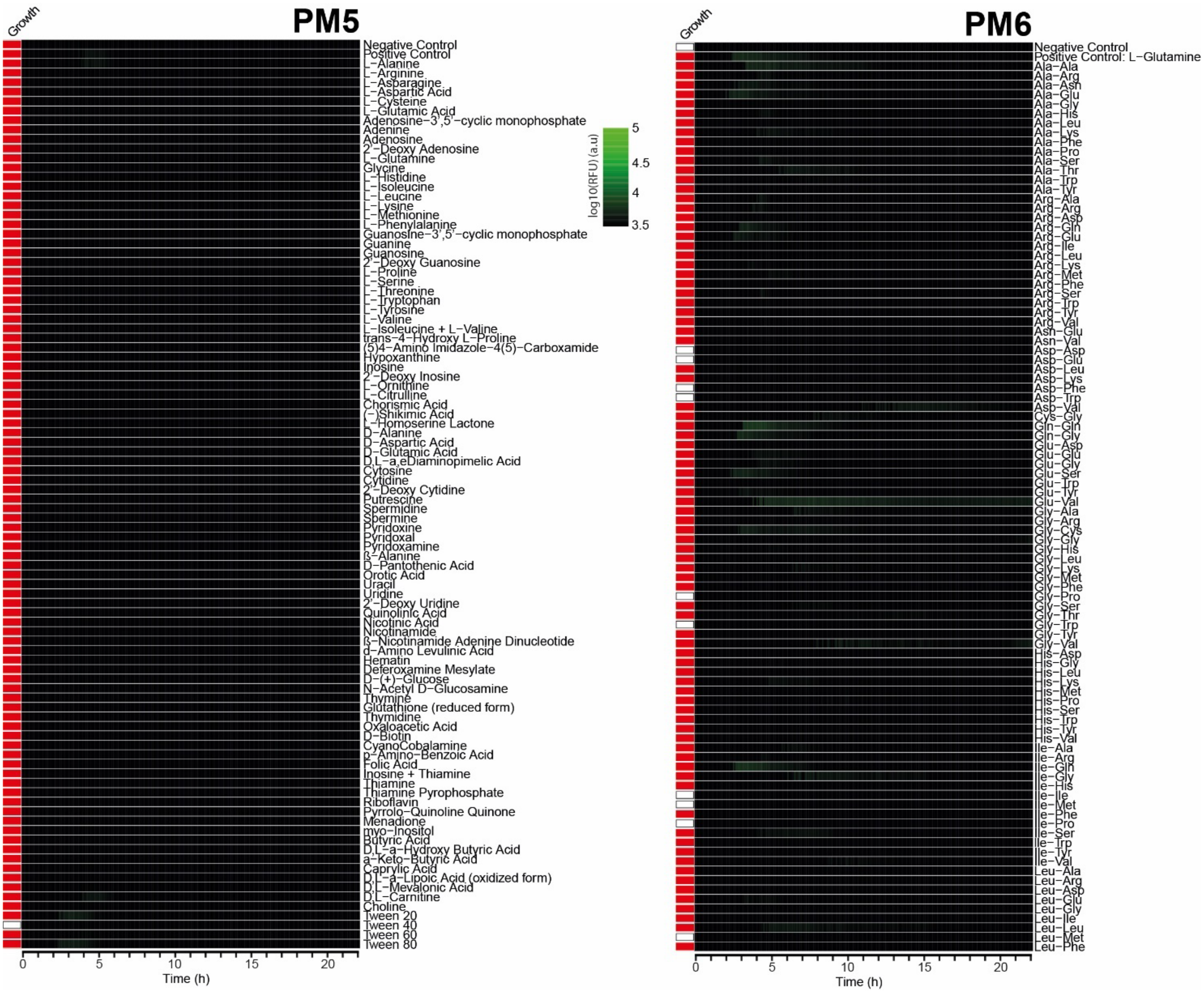


**Figure S3: Monitoring reporter fusion induction of R-tailocin expression in different biosynthetic pathways (PM5) and under different peptide compositions (PM6).**

Cells of *P. protegens* CHA0 carrying the R-tailocin reporter fusion were inoculated into Biolog PM5 and PM6 plates, each well containing a distinct nitrogen source. The inoculation medium consisted of Minimal Medium (MM) without any added nitrogen source for PM6, ensuring that the compounds provided in the Biolog plate served as the sole available nitrogen sources for bacterial growth. RFU (Relative Fluorescence Unit) represents the fluorescence signal normalized by optical density (OD). Red squares indicate wells where bacterial growth occurred, while white squares indicate wells with no detectable growth.

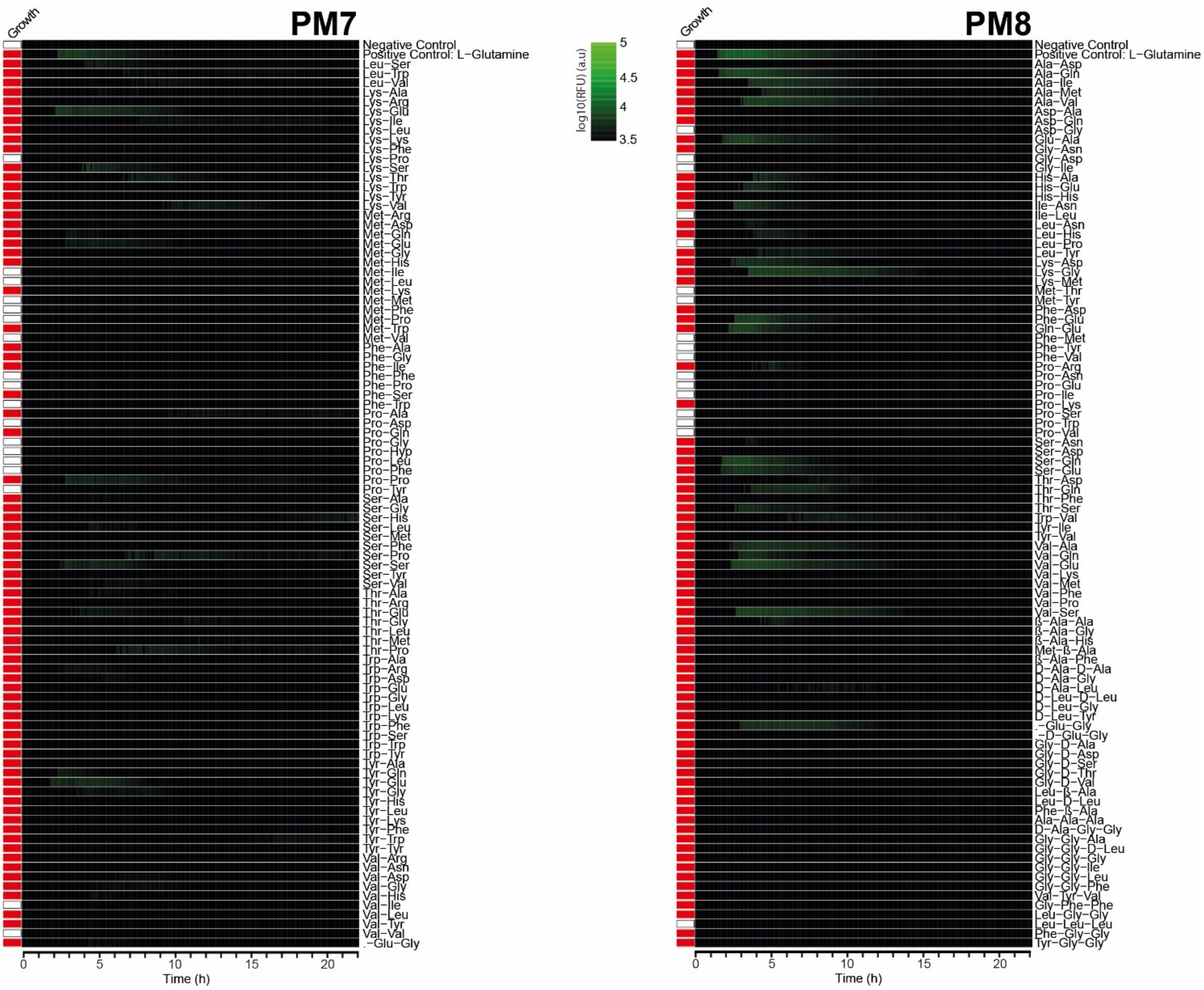


**Figure S4: Monitoring reporter fusion induction of R-tailocin expression under different peptide compositions (PM7 and PM8).**

Cells of *P. protegens* CHA0 carrying the R-tailocin reporter fusion were inoculated into Biolog PM7 and PM8 plates, each containing a distinct nitrogen source. The inoculation medium consisted of Minimal Medium (MM) without any added nitrogen source, ensuring that the compounds provided in the Biolog plates served as the sole available nitrogen sources for bacterial growth. RFU (Relative Fluorescence Unit) represents the fluorescence signal normalized by optical density (OD). Red squares indicate wells where bacterial growth occurred, while white squares indicate wells with no detectable growth.

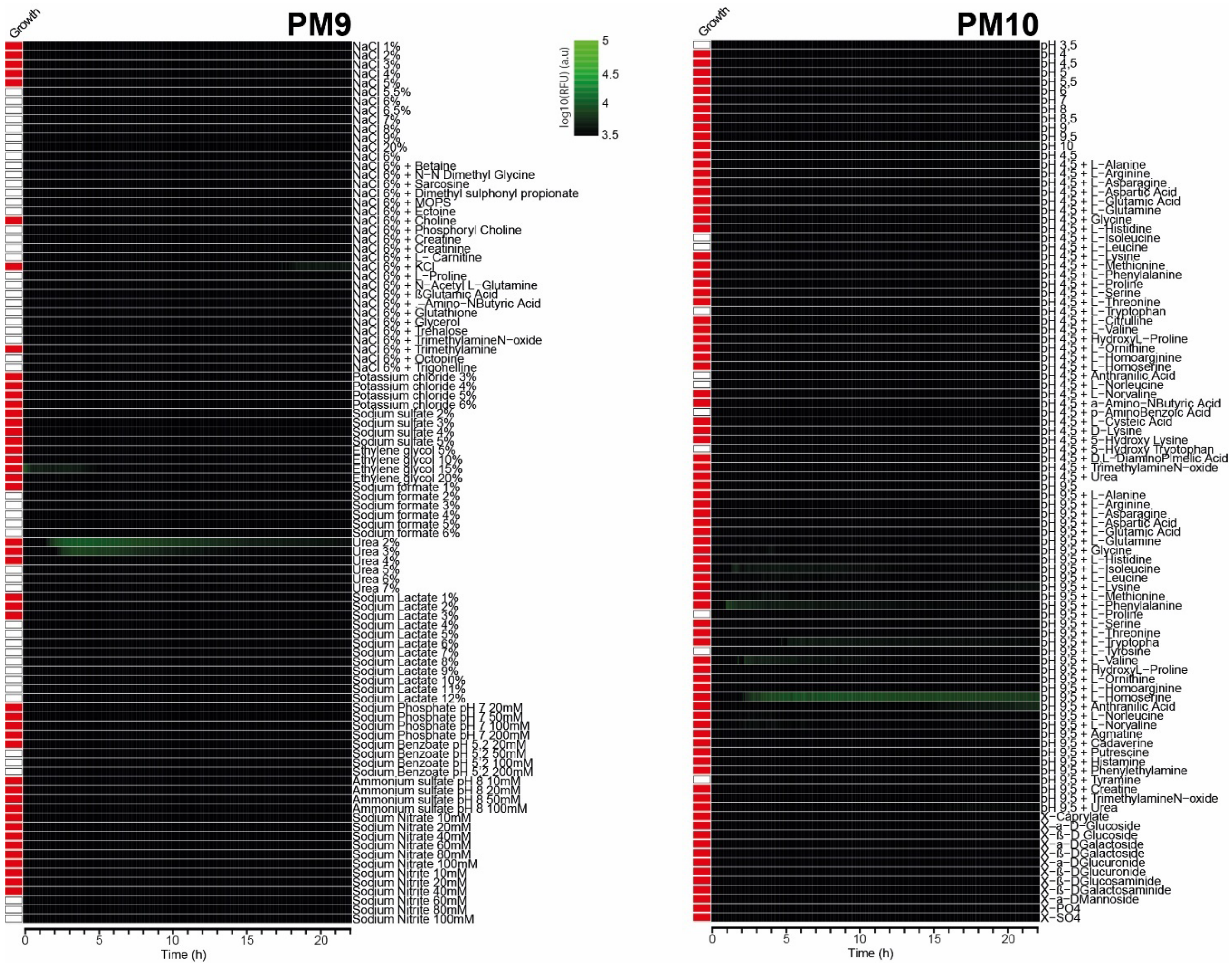


**Figure S5: Monitoring reporter fusion induction of R-tailocin expression under different osmotic stress (PM9) and pH (PM10) conditions.**

The inoculation medium consisted of Minimal Medium (MM) supplemented with glucose (80 mM) as the carbon source, ensuring that observed changes in fluorescence were primarily due to the tested conditions rather than nutrient limitation. RFU (Relative Fluorescence Unit) represents the fluorescence signal normalized by optical density (OD). Red squares indicate wells where bacterial growth occurred, while white squares indicate wells with no detectable growth.

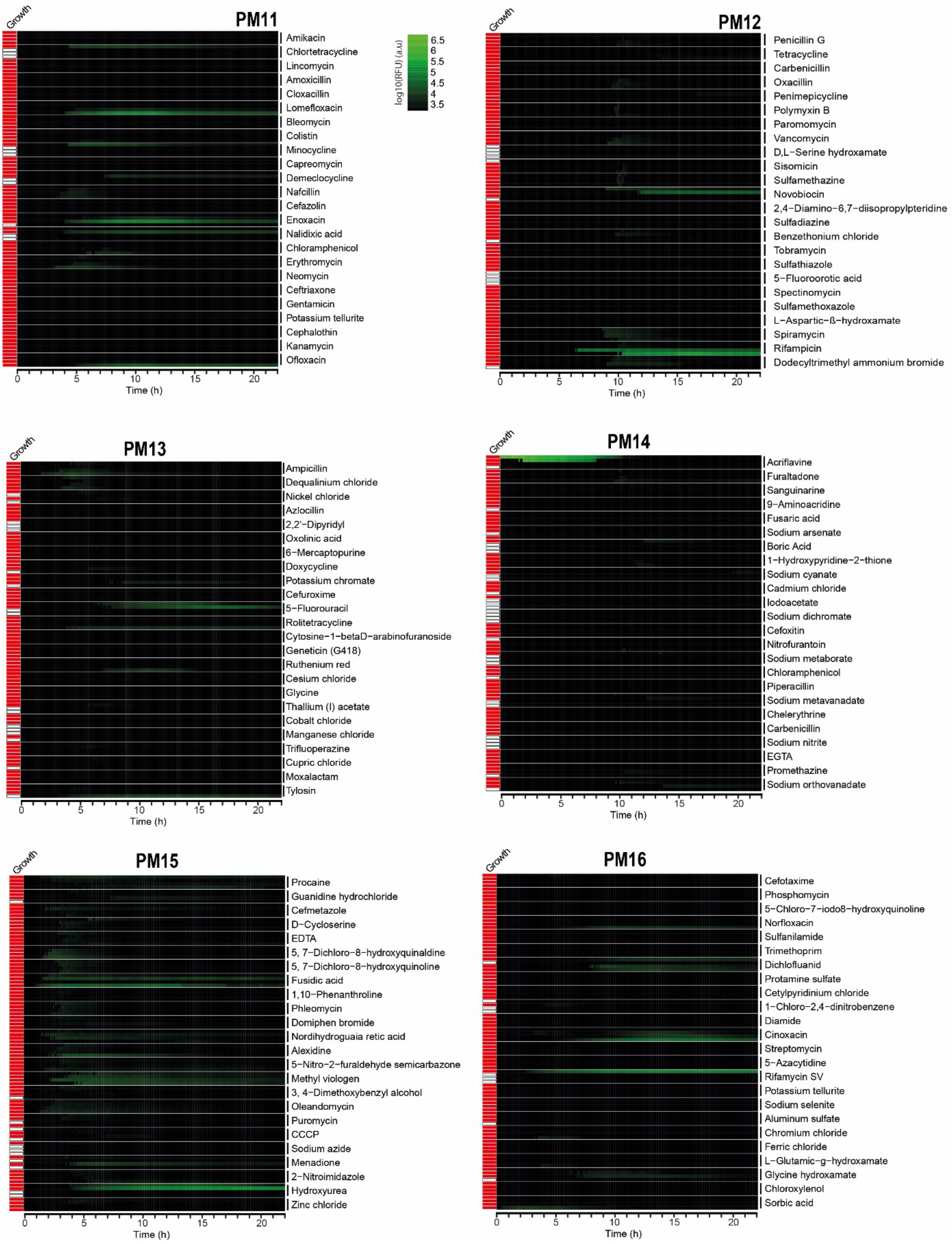


**Figure S6: Monitoring reporter fusion induction of R-tailocin expression under different chemical stress (1/2).**

The inoculation medium consisted of Minimal Medium (MM) supplemented with glucose (80 mM) as the carbon source, ensuring that observed changes in fluorescence were primarily due to the tested compounds rather than nutrient limitation. RFU (Relative Fluorescence Unit) represents the fluorescence signal normalized by optical density (OD). Red squares indicate wells where bacterial growth occurred, while white squares indicate wells with no detectable growth. Four increasing concentrations were tested for each compound.

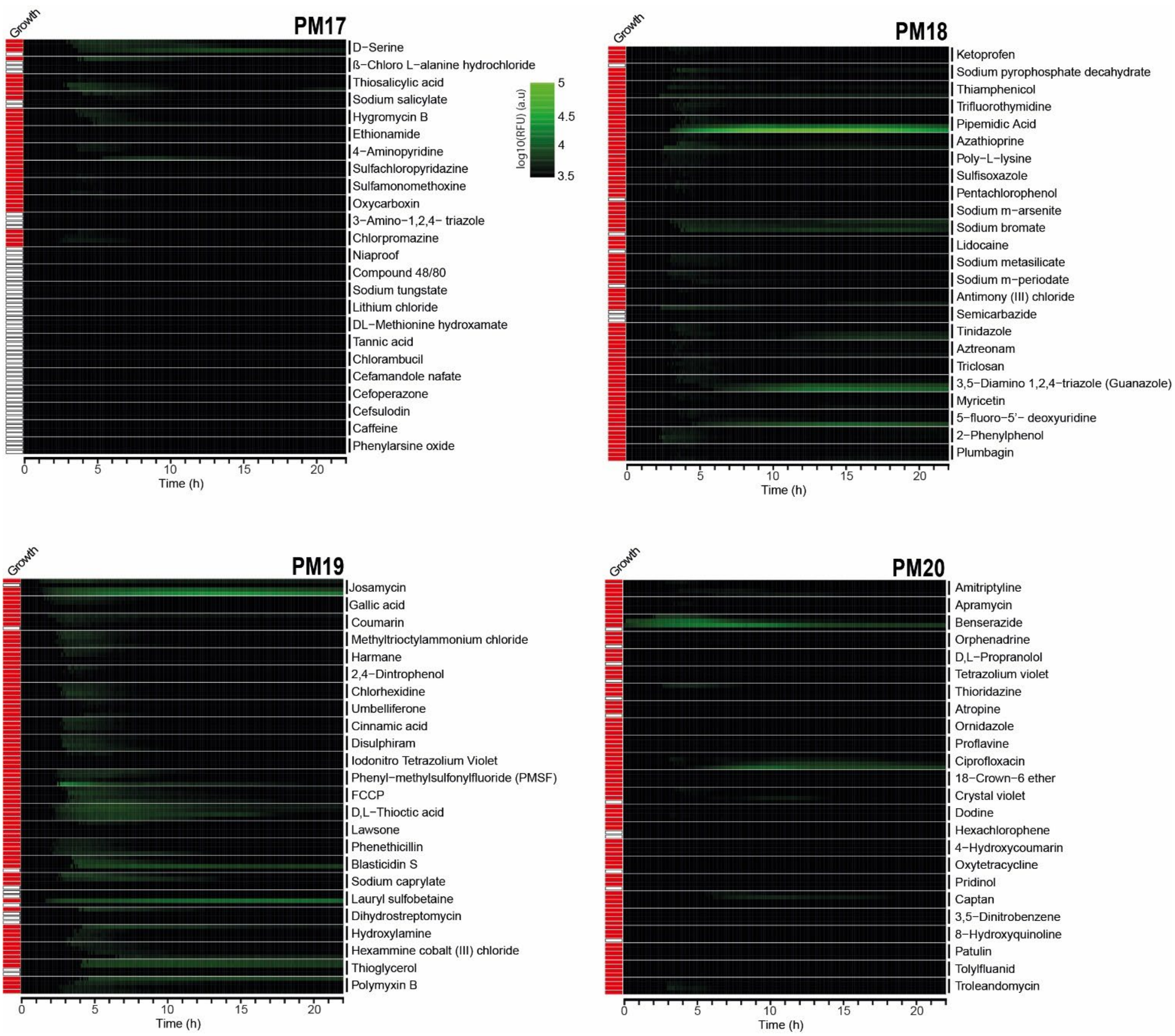


**Figure S7: Monitoring reporter fusion induction of R-tailocin expression under different chemical stress (2/2).**

The inoculation medium consisted of Minimal Medium (MM) supplemented with glucose (80 mM) as the carbon source, ensuring that observed changes in fluorescence were primarily due to the tested compounds rather than nutrient limitation. RFU (Relative Fluorescence Unit) represents the fluorescence signal normalized by optical density (OD). Red squares indicate wells where bacterial growth occurred, while white squares indicate wells with no detectable growth. Four increasing concentrations were tested for each compound.
